# Highly efficient bio-catalytic oxygen reduction coupled to long-range electron transport in cable bacteria

**DOI:** 10.64898/2026.01.20.700514

**Authors:** Dmitrii Pankratov, Anwar Hiralal, Silvia Hidalgo Martinez, Jeanine S. Geelhoed, Filip J. R. Meysman

## Abstract

Multicellular cable bacteria are capable of transferring electrons over centimeter distances through an internal array of conductive fibers. These long, filamentous bacteria function as a living electrochemical cell, performing sulfide reduction at one end and oxygen reduction at the other end. To investigate how O_2_ reduction is linked to the long-distance electron transport along the conductive fibers, we performed a detailed electrochemical characterization of native filaments as well as extracted “fiber skeletons” without membranes or cytoplasm. Our data show that fibers skeletons only perform longitudinal electron transport and are not electrochemically active towards oxygen. This opposes a previous proposition that the conductive fiber network displays electrocatalytic behavior towards oxygen. Still, native cable bacterium filaments are capable of high oxygen reduction rates, thus demonstrating that dedicated enzyme systems in the periplasm or inner membrane are responsible for O_2_ reduction. Together, our data provide empirical support for a model in which diffusible c-type cytochromes mediate electron transport through the periplasm, shuttling electrons between separate respiratory complexes and the conductive fiber network. As such, our study resolves a crucial aspect of the unique electrogenic metabolism in cable bacteria, and clarifies the application potential of the highly conductive fibers in Bio-electrochemical System technologies.

## 1. INTRODUCTION

Cable bacteria are multicellular bacteria that thrive globally in freshwater and marine sediments [1–4]. They stand out in the microbial world due to their unique, electrogenic metabolism, in which electrons are transported over centimeter-scale distances [5, 6]. This long-distance electron transport electrically couples the oxidation of sulfide in deeper sediment layers to the reduction of oxygen or nitrate near the sediment surface [1, 7], and this capability equips cable bacteria with a competitive advantage over other single-celled sulfur-oxidizing bacteria [6]. Cable bacteria consist of linear filaments encompassing up to thousands of cells. On the outer surface, they display a set of parallel ridges, each of which embeds a conductive fiber that runs along the whole length of the filament [8]. Long-distance electron transport is guided through this highly ordered network of conductive fibers [8–10]. Detailed electrical characterization has shown that the fiber conductivity is higher than that of most semiconductor materials used in electronics [10–12], reaching values up to 564 S cm^-1^ [13]. Such conductivity is remarkable for biological material [10], and is thought to be linked to an elaborate supramolecular nickel-organic framework structure essential for conductance [14, 15].

Electrons are “uploaded” to or “downloaded” from the fiber network by the process of electrogenic sulfur oxidation, which can be depicted as a spatial separation of two redox half-reactions, inducing a clear division of labor between two types of cells [16]. In deeper sediment, the so-called “anodic” cells of the cable bacteria filaments perform the oxidation half-reaction of sulfide, which is locally produced by sulfate reduction or iron-sulfide dissolution [17]:

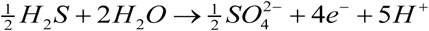

Near the sediment-water interface, the so-called “cathodic” cells perform reduction half-reaction of oxygen, which diffuses from the overlying water into the sediment and is rapidly consumed within the first few millimeters:

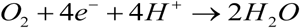

Apart from O_2_, cable bacteria can utilize nitrate as an alternative electron acceptor, though when both are present, O_2_ seems to be preferred [7]. De facto, cable bacteria function as a living electrochemical cell, driven by the physically separated redox half-reactions of oxygen reduction and sulfide oxidation, while the conductive fiber network acts like a power line network that ensures the electron transport from the anodic to the cathodic cells in the filament.

The mechanism of oxygen reduction and its link to long-range electron transport is the subject of ongoing debate, as there is conflicting evidence on how the mechanism of electron transport exactly functions (i.e. the interplay between uploading, transport and downloading of electrons) [18, 19]. In one model, the conductive fibers exclusively possess an electron transport function, and lack oxygen reduction activity [18]. In this view, anodic sulfide oxidation and cathodic oxygen reduction are catalyzed by dedicated enzyme systems, embedded in either the periplasm or integrated within the inner membrane [18]. In contrast, other studies suggest that the fiber network itself is capable of catalysis, and so in this model, oxygen reduction occurs directly on the fibers and is not mediated by membrane-bound enzymes [19]. Here, we investigated the electrocatalytic properties of two types of cable bacterium filaments (native and chemically extracted) using cyclic voltammetry and electrochemical gating. Our results provide a new empirically supported model for the link between oxygen reduction and long-range electron transport.

## 2. EXPERIMENTAL

### 2.1 Materials

All chemicals were of at least analytical grade and used without further purification. Na_2_HPO_4_⋅2H_2_O, NaH_2_PO_4_⋅H_2_O, H_2_SO_4_, 6-mercapto-1-hexanol, N-acetyl-L-methionine were purchased from Sigma-Aldrich. All solutions were prepared using deionized water (18.2 MΩ cm) produced with an Arium system (Sartorius, Germany).

### 2.2 Culturing conditions

Both cable bacteria species used in this study were originally sampled from Rattekaai salt marsh (the Netherlands; 51.4391°N, 4.1697°E). For *Ca*. E. gigas, sediment was sieved, homogenized by mixing and repacked into transparent PVC core liner tubes (36 mm diameter, 100 mm height) closed at the bottom with a rubber stopper. The sediment cores were incubated in the dark in artificial seawater (Instant Ocean Sea Salt, salinity 30, 20°C) under continuous aeration for several weeks. For *Ca*. E. scaldis GW3-3, a continuation of the original clonal enrichment culture was used, which has been described in [26]. For both enrichment cultures, incubation lasted several weeks until the typical geochemical fingerprint of cable bacteria was detected [62]. Modification with N-acetyl-L-methionine was performed for the sediment core with actively growing populations, similar to the earlier developed and optimized procedure of isotope labeling of cable bacteria [14]. 500 μL of 100 mM N-acetyl-L-methionine solution was injected using a 50 μL gas chromatography syringe into each of the 3 “subcores” formed by the plastic cylinders (10 mm diameter, 50 mm height) inserted into the sediment to a depth of approximately 45 mm. After the 4-day incubation cable bacteria were harvested from the sediment.

Individual cable bacterium filaments were picked under a binocular microscope with fine glass hooks custom-made from Pasteur pipettes and subsequently washed in several droplets of deionized water.

### 2.3 Preparation of interdigitated electrodes

Interdigitated gold electrodes (IGEs, 125×2 digits, bands/gaps - 10 µm) were purchased from Metrohm DropSens (Spain). IGEs were treated by electrochemical cycling in 0.5 M H_2_SO_4_ from −0.2 to 1.7 V vs saturated calomel electrode (SCE) with a scan rate of 0.1 V s^−1^ until steady-state voltammograms were obtained. The electrodes were rinsed with deionized water, incubated for 24 h in an 8 mM solution of 6-mercapto-1-hexanol, washed with water and dried.

### 2.4 Instrumentation and measurements

Electrochemical measurements were performed using a PalmSens3 bipotentiostat (for electrochemical gating) and PalmSens4 potentiostat (Palmsens BV, Houten, The Netherlands), controlled by the PSTrace software. In electrochemical gating, two parts of IGEs were connected separately as two working electrodes and were used as the source and the drain. In voltammetric experiments, two parts of IGEs were shortcut by a strip of copper tape, and the whole gold surface was utilized as a working electrode. A glassy carbon rod and SCE were used as auxiliary and reference electrodes. Cyclic voltammograms were recorded on IGEs at a 20 mV s^-1^ scan rate in a 50 mM potassium phosphate buffer solution (pH 7.0). The onset potential was determined as the potential at which bioelectrocatalytic current exceeds a 3-fold deviation of the noise level from currents in the deoxygenated electrolyte [63].

### 2.5 Microscopy

Light microscopy images were obtained using a Zeiss Axioplan 2 microscope equipped with an Exi Blue Camera (QImaging). The length and width of the filaments were measured using Image Pro Insight (Media Cybernetics). Postprocessing of the images was done using Fiji software [64]. Atomic force microscopy (AFM) was performed using an XE-100 AFM system (Park Systems) operating in tapping mode, using an aluminum SPM probe with a tip radius of <10 nm (AppNano ACTA-200) and with a nominal spring constant of 13–77 N/m. Topography and amplitude images were recorded and processed with the software Gwyddion [65].

## 3. RESULTS

### 3.1 Cyclic voltammetry of fiber skeletons

The oxygen reduction activity was assessed for cable bacterium filaments belonging to the *Ca.* Electrothrix gigas species, which is marked by a large diameter (> 2.5 microns) and a high number of outer surface ridges (> 50) [20]. Sequential extraction of native filaments removes the lipid membranes and cytoplasm, thus producing a so-called “fiber skeleton”, which retains the conductive fiber network connected by an underlying polysaccharide sheath [8]. Fiber skeletons retain a similar conductivity as native filaments [10], thus indicating that the extraction procedure does not affect the structure and transport properties of the conductive fiber network. However, conflicting evidence exists about the capability of these fiber skeletons to perform O_2_ reduction. Previous reports claim to observe oxygen reduction activity [19], while in other experiments, such activity was not detected [10]. To investigate the electrocatalytic behavior, we deposited fiber skeletons onto interdigitated gold electrodes (IGEs) and performed cyclic voltammetry (CV; Figure 1A – left configuration). Initially, the electrolyte solution remained anoxic, after which we let the oxygen level gradually increase. The recorded CVs do not show any dependence on the O_2_ concentration of the electrolyte medium (50 mM potassium phosphate buffer solution, pH 7.0, T = 23°C) (Figure 1C). Fiber skeletons hence do not reveal any oxygen reduction activity, thus confirming the results reported in [10] and contradicting the claim that the fibers can catalyze the reversible interconversion of oxygen and water [19].

**Figure 1.**
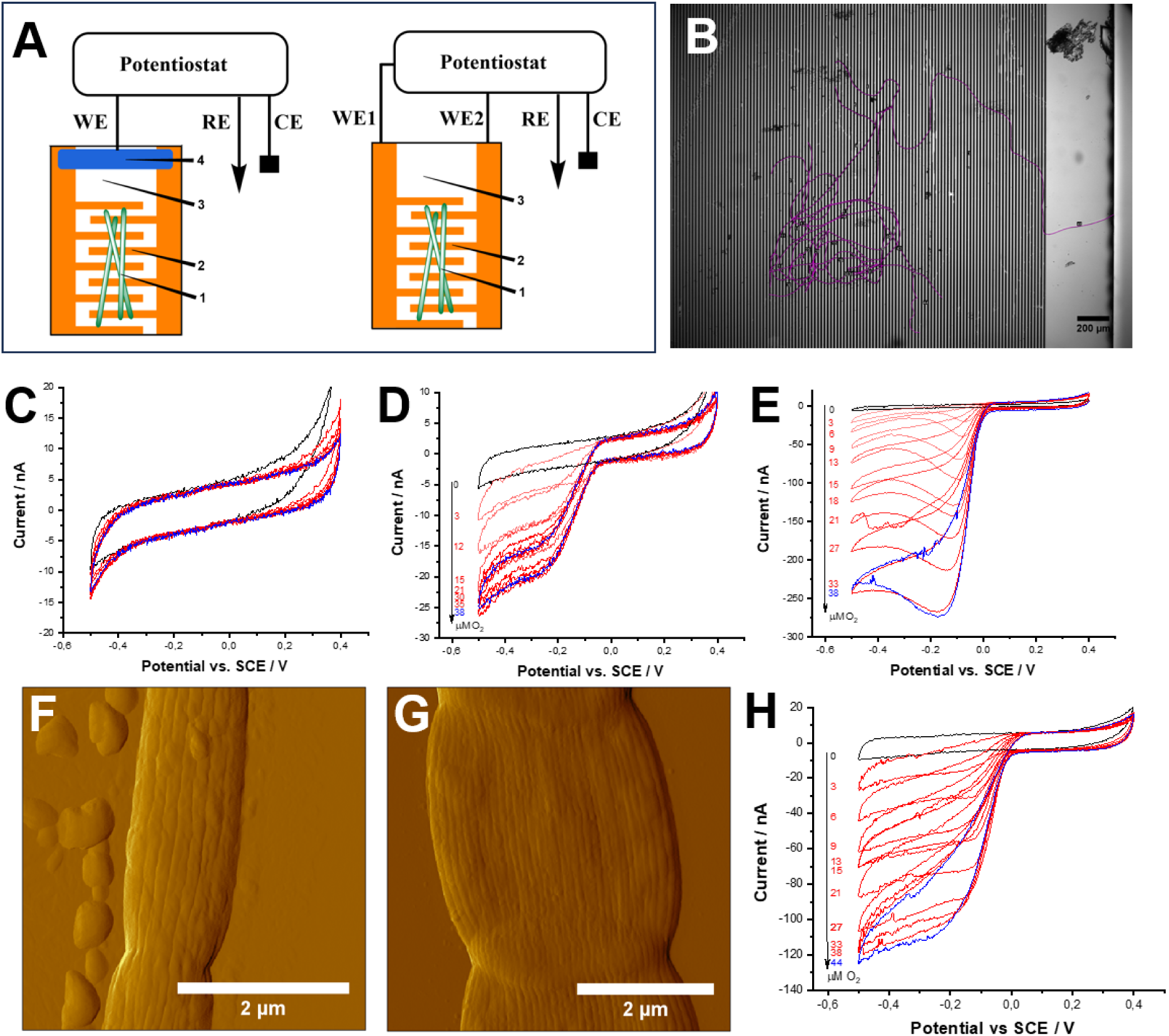
Electrochemical characterization of cable bacterium filaments. (A) Schematic of electrochemical setup used for cyclic voltammetry (left) and electrochemical gating (right): WE – working electrode (WE1 and WE2 – source and drain in electrochemical gating experiments, respectively), CE – counter electrode, RE – reference electrode, 1 – intact cable bacteria, 2 – gold surface, 3 – transparent support, 4 – strip of copper tape. (B) Representative micrograph of a bundle of intact cable bacteria deposited on the IGE. Overlying lines (purple) are drawn to determine the total length of the filaments. (C-E, H) Representative cyclic voltammograms of three different types of filaments as a function of the added O_2_ concentration. (C) Extracted fiber skeletons do not show any reactivity towards O_2_ in the range of 0 (black) – 47 (blue) µM O_2_. Red CVs correspond to intermediate O_2_ concentrations. (D) Native filaments with low O_2_ reduction activity. (E) Native filaments with high O_2_ reduction activity. (F-G) AFM images of native filaments of *Ca*. E. scaldis (F) and *Ca*. E. gigas (G). (H) Representative CVs of native filaments from *Ca*. E. scaldis at different concentrations of O_2_. All scans were recorded in a phosphate buffer (50 mM, pH 7.0; T=23°C) at a scan rate of 20 mV s^-1^.

### 3.2 Cellular-specific oxygen reduction rates in native filaments

Cable bacteria display among the highest cell-specific oxygen consumption rates within the microbial realm [16, 21], and so it is crucially important to understand how such high rates can be achieved. To this end, cyclic voltammograms were additionally recorded on IGEs for native filaments of *Ca.* E. gigas, in a similar way as was done for fiber skeletons (Figure 1A – left configuration). The resulting CVs show the development of a distinct catalytic wave when the O_2_ concentration in the medium increases (Figure 1D,E). The onset potential of the catalytic wave occurs within the +20-50 mV range versus saturated calomel electrode (SCE) at the experimental conditions investigated (T=23°C, pH 7). This onset potential is considerably higher than typically recorded for other microorganisms capable of oxygen reduction (ca. −170 mV) [22], indicating that less energy can be theoretically gained from cathodic O_2_ reduction, but still comprises a significant overpotential compared to the redox equilibrium potential of the O_2_/H_2_O couple under the same conditions (+580 mV).

When recording CVs of replicate cable bacteria samples (n=7), we noticed two distinct types of current-voltage responses. In one set of samples, referred to as low-rate filaments (LRF), we saw a gradual increase in the bioelectrocatalytic current when the potential was decreased (Figure 1D). In a second set of samples, referred to as high-rate filaments (HRF), the cathodic current increased rapidly, until a maximum was reached, after which the current decreased again (Figure 1E). The latter likely reflects a transport limitation related to the oxygen supply towards the planar, non-rotating IGE. Local depletion of O_2_ in an electrical double layer leads to a decrease in electrocatalytic current [23]. The potential at which the current started to decrease was dependent on the O_2_ level in the electrolyte, shifting from −70 to −170 mV when the O_2_ concentration in the medium was increased from 3 to 33 µM (Figure 1C). In both LRF and HRF samples, the maximum cathodic current was reached at *ca*. 35 μM O_2_, which is slightly lower than found in our previous work (50 μM O_2_) [16]. This may be related to the more precise control of the O_2_ concentration in the buffer solution here (which was improved compared to the previous setup). In the absence of cable bacteria (control electrode with a bare thiolated gold surface), the cathodic current was not influenced by the O_2_ concentration over the potential range examined (Figure S1). This indicates that the observed O_2_ reduction activity is induced by catalytic components embedded in native cable bacteria, but located outside of the fiber network (as fiber skeletons remain non-responsive to O_2_, Figure 1C).

For each sample, the total length of filaments deposited on the electrode was quantified by light microscopy and ensuing image analysis (19.2–77.4 mm, Figure 1B), enabling the calculation of cellular O_2_ reduction rates from the observed maximum electrocatalytic current (Table S1). The different samples showed a wide variation in the number of cells in contact with the electrode surface (3,200-12,900 cells), in addition to a sizeable variation in cellular O_2_ reduction rates (14-130×10^-18^ mol O_2_ cell^−1^ s^−1^). In general, LRF samples without diffusion limitation showed a lower cellular O_2_ reduction rate than HRF samples with diffusion limitation.

### 3.3 Dependence of the oxygen reduction activity on O_2_ and pH

The maximum current attained (as derived from the CVs) can be plotted as a function of the O_2_ concentration in the electrolyte buffer (Figure 2A,D). Despite the notable difference in the shape of the CVs, LRF and HRF samples were characterized by the same apparent Michaelis-Menten constant (*K’_m_*), viz. 27.5 ± 4.3 and 27.5 ± 15.0 µM, respectively (Table S2, Figure 2A,D). Since *K’_m_* values are not affected by the concentration of the enzyme, this suggests that LRF and HRF operate a similar enzymatic machinery for O_2_ reduction. Accordingly, the observed variation in cellular O_2_ reduction rates (Table S1) must be related to the concentration of biocatalyst within the filaments.

**Figure 2.**
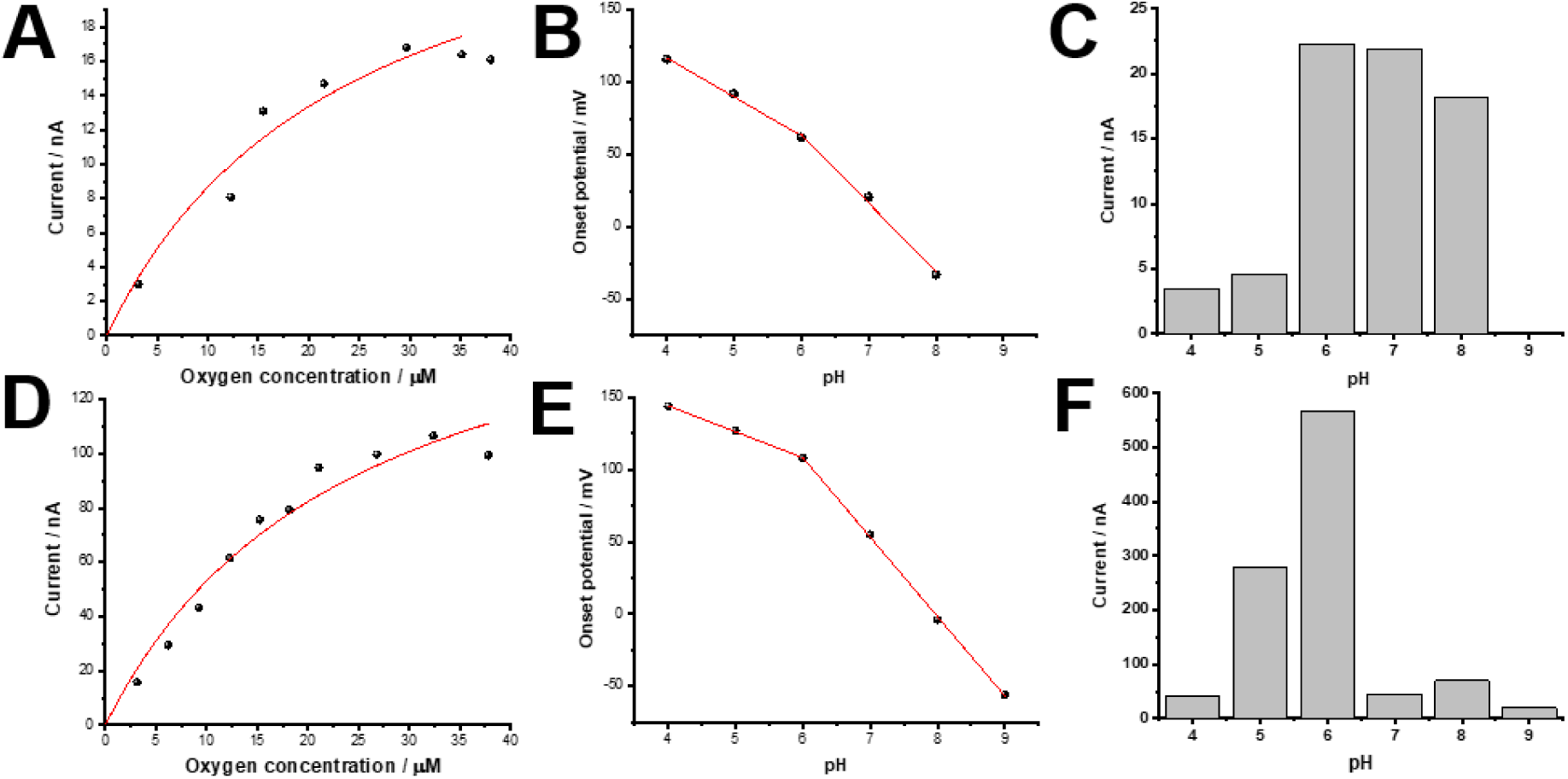
Oxygen reduction activity of two types of cable bacteria: low-rate filaments (LRF- top row panels A-C), and high-rate filaments (LRF- top row panels A-C). (A, D) Representative current versus [O_2_] plots. (B,E) pH dependence of the onset potential of the oxygen reduction rate. (C,F) Bioelectrocatalytic current as a function of pH.

This hypothesis is supported by the pH dependence of the onset potential of O_2_ reduction, which is similar for the LRF and HRF samples and shows a marked change in the slope around pH 6 (Figure 2B,E). Above pH 6, the onset potential of LRF and HRF samples decreases with ca. −54 mV per pH unit, formally corresponding to a 1e^−^/1H^+^ process. In the region below pH 6, the slope is only about half, indicating a 2e^−^/1H^+^ mechanism. In contrast, the pH dependence of the maximum current for O_2_ reduction reveals a difference between the LRF and HRF samples (Figure 2C,F). For LRF, this current reaches a maximum at pH 6 and remains constant afterward (Figure 2C, S2). For HRF, the current also attains a maximum at pH 6, but subsequently, the current notably drops when the pH increases to 7 or higher (Figure 2F, S2).

The CVs above were all recorded at low O_2_ concentrations, at which the O_2_ reduction rate remained constant during the measurement. Yet, when the buffer solution was saturated with air, the O_2_ reduction rate decreased rapidly with time (Figure S3). The half-inactivation time, defined as the period over which the O_2_ reduction rate decreased to half its original value, amounted to 0.24 and 1.2 hours for the LRF and HRF at O_2_-saturated conditions, respectively. The decrease of the O_2_ reduction rate was slower for HRF filaments, and so apparently, a more rapid oxygen consumption by HRF leads to a longer inactivation time.

Filaments grown in the presence of 100 mM N-acetyl-L-methionine did not show any dependence on the O_2_ concentration at CVs (Figure S4). N-acetyl-L-methionine is known to deactivate c-type cytochromes by changing their redox potential through heme ligation [24, 25]. As a result, the deactivation of c-type cytochromes appears to suppress the O_2_ reduction activity.

### 3.4 O_2_ reduction activity in different cable bacterium species

The above data were all recorded for filaments belonging to the *Ca.* E. gigas species (Figure 1G). To investigate whether the same response is also seen in other cable bacteria, cyclic voltammetry was performed on native filaments from a different species of cable bacteria, *Ca*. E. scaldis GW3-3 [26], which has a very different morphology (Figure 1F). Whereas *Ca*. E. scaldis GW3-3 is relatively small in diameter (width = 0.75±0.09 µm) and has fewer ridges (∼12 ridges; *n* = 8) [13], *Ca.* E. gigas is much wider (width ranging from 2.5–8 µm) and has over 5 times more ridges (>52) [8, 20]. Secondly, both species exhibit a (partially) different genetic repertoire for O_2_ reduction. Both species encode a periplasmic truncated hemoglobin postulated to have major O_2_ reduction capacity, as well as a CydA homolog of the cytochrome *bd* quinol oxidase complex and periplasmic cytochromes that could potentially reduce O_2_ [13, 18]. Yet in addition, *Ca*. E. scaldis also encodes the membrane-bound cytochrome *c* oxidase complex CoxABCD [13], which is completely absent in *Ca*. E. gigas species [13, 18], and may function as an O_2_ reductase. As can be seen in Figure 1H, the voltammetric behavior of *Ca*. E. scaldis is identical to that of *Ca*. E. gigas, despite the differences in morphology and genetic repertoire. The onset potential of ORR (*ca*. 30 mV), the limiting O_2_ concentration of substrate (35 µM O_2_) and the value of *K’_m_* (27 µM, Table S2) are similar to *Ca*. E. gigas. This suggests that the putative presence of an additional oxygen-reducing enzyme (cytochrome *c* oxidase complex CoxABCD) in *Ca*. E. scaldis GW3-3 does not noticeably change the affinity for O_2_ reduction (although we don’t know to what extent the protein complex is effectively expressed under the conditions investigated).

### 3.5 Relation between O_2_ reduction and long-range conductivity

To further characterize the redox sites, we applied the electrochemical gating technique [13, 27] to the clumps of native filaments deposited onto IGE arrays submerged in an oxygen-free phosphate buffer solution (Fig. 1A – right panel; see also Materials and methods). By conjointly changing the potential of the source and drain electrodes, one measures the conductance as a function of the oxidation state of the redox moieties embedded in the conductive filament. The source and drain potentials (E_S_ and E_D_) are controlled by a bipotentiostat and scanned simultaneously at the same rate ν, while maintaining a constant small source-drain voltage bias (V_SD_ = E_D_ − E_S_).

Electrochemical gating of native filaments showed a characteristic dome-shaped dependence of the source-drain current on the source electrode potential (Figure 3). This response is markedly different from the previously recorded electrochemical gating behavior of fiber skeletons, in which the gating current remains independent of the gating potential [13]. The dome-shaped dependence indicates that redox-active charge carriers must be present within the conduction path between the source and drain electrodes, similar to the gating behavior reported for cells and biofilms of *Shewanella oneidensis* MR-1 [28], *Geobacter sulfurreducens* [29, 30] or a mixed *Marinobacter-Chromatiaceae-Labrenzia* community [31]. The observed potential at the maximum gating current (*ca*. −100 mV) corresponds to the formal potential of the redox-active charge carriers in native cable bacterium filaments [27].

**Figure 3.**
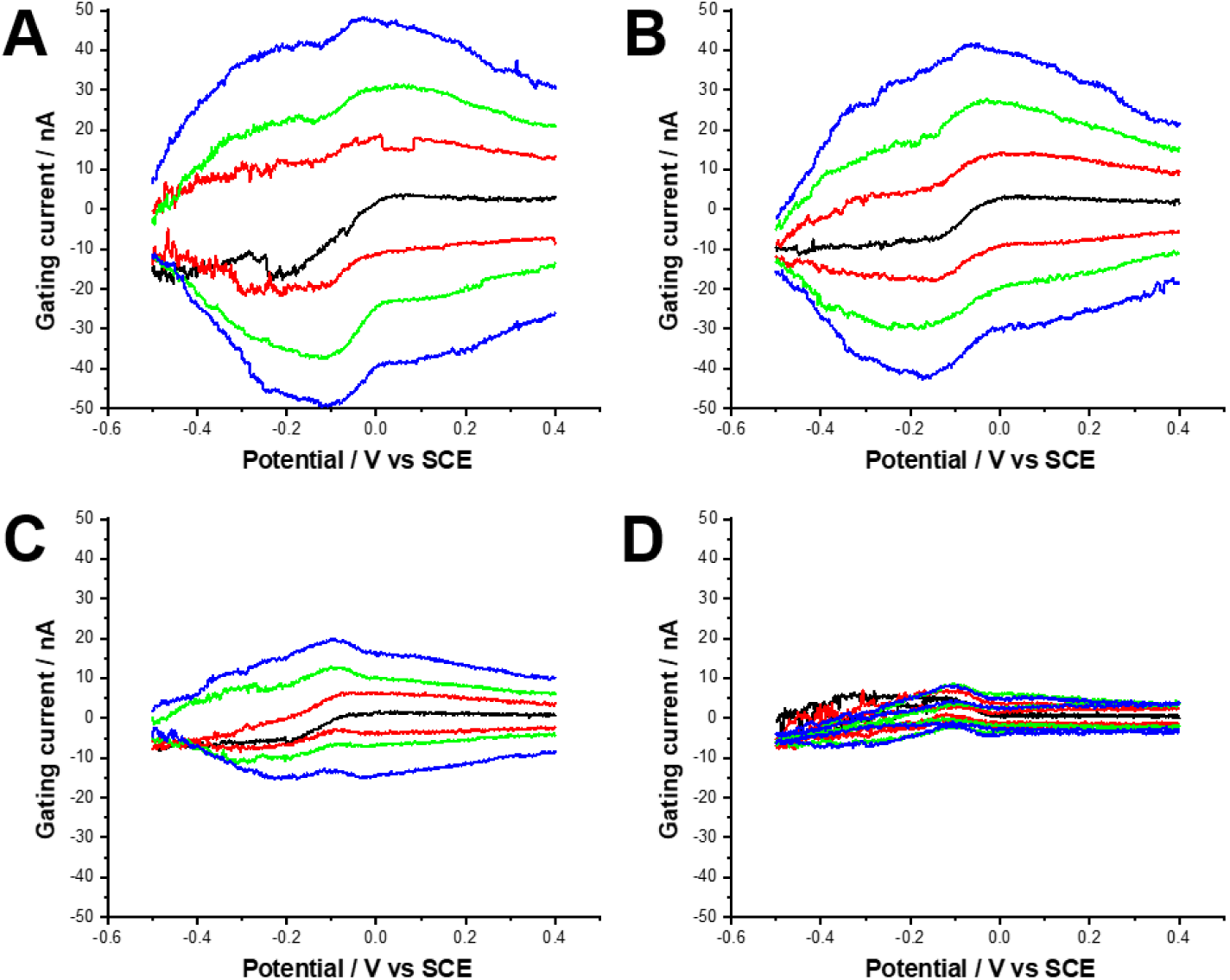
Dependence of source-drain current on the source electrode potential for intact cable bacteria in phosphate buffer (pH 7) at different concentrations of oxygen: 0 µM (A), 3.2 µM (B), 9.5 µM (C), 21.8 µM (D). Source-drain voltage: 0 mV (black curves), 3 mV (red curves), 6 mV (green curves), and 10 mV (blue curves). Scan rate: 5 mV s^-1^. Background current output was recorded from bare thiolated IGEs under the same conditions and subtracted from the curves presented.

The gating current decreased when O_2_ was added to the electrolyte solution, until conductivity completely vanished when the buffer became air-saturated (Figure 3, Figure S5). Afterwards, CV scans revealed that these filaments, exposed to aerobic conditions and lacking conductivity, retained their O_2_ reduction activity (Figure S5). This suggests that the presence of oxygen rapidly caused a severe degradation of fiber conductivity, but that it does not affect the activity of the O_2_ reduction system to the same extent (as it shows much slower performance degradation). This confirms that activity of fibers and O_2_ reduction complexes is distinct, as oxygen reduction in native cable bacteria filaments remains possible even in the absence of fiber conductivity.

## 4. DISCUSSION

Cable bacteria perform long-range conduction in which electrons are first harvested from anodic sulfide reduction in deeper sediment layers, and then transported along the intern fiber network up towards the oxic zone, where O_2_ reduction takes place [1, 10]. Given that cable bacteria typically have 10 times less cells in the oxic zone compared to the suboxic zone [32], this implies these “oxic” cells must process a high flow of electrons, and so they should embed a highly efficient O_2_ reduction machinery. It has been hypothesised that O_2_ reduction does not conserve energy, but simply serves to channel a metabolic waste product (i.e. electrons) to the external environment as efficiently as possible, for the greater good of the multicellular organism [16]. The high onset potential for oxygen reduction as observed here (ca. −170 mV) relative to other O_2_-consuming bacteria [22] supports this idea. Our results provide insights into how this putative “altruistic” O_2_ reduction machinery functions, and how it is linked to the electron transport along the periplasmic fiber network.

### 4.1 The fiber network does not show oxygen reduction activity

Fiber skeletons do not reveal any oxygen reduction activity (Figure 1C). As a consequence, the function of the fiber network inside the cable bacteria is to enable long-distance electron transport, while oxygen reduction must be catalyzed by enzymes external to the fiber network. This finding is consistent with previous experiments [10], but in conflict with the results presented in [19], which put forward that fiber skeletons do show O_2_ reduction activity. These conflicting observations may possibly be reconciled by incomplete extraction of fiber skeletons. When cable bacterium filaments are only partially extracted, enzymes capable of oxygen reduction (e.g. cytochromes) can remain attached to the fiber skeletons, and this may induce catalytic activity as seen in [19]. Therefore, it is adamant to verify the outcome and quality of the fiber skeleton extraction procedure, which is possible by Raman microscopy [9, 10, 14]. Fiber skeletons display a characteristic Raman fingerprint, which originates from the Ni/Si metal-organic framework that is embedded in the fibers and which enables the long-range electron transport [9]. Additionally, fiber skeletons should not show any of the strong resonant peaks of cytochromes, which are removed during the SDS-EDTA extraction [14]. As such, the presence of resonant cytochrome peaks hence provides an indicator of incomplete extraction of fiber skeletons.

### 4.2 Native filaments show high potential cellular-specific oxygen reduction rates

In contrast to fiber skeletons, native filaments of cable bacteria do reveal high O_2_ reduction activity (Figure 1D,E). Foremost, our data show cellular O_2_ reduction rates ranging between 14 and 130×10^-18^ mol O_2_ cell^−1^ s^−1^ which confirms previous observations that cell-specific oxygen consumption rates in cable bacteria are high [16]. The maximum value observed is ∼2-fold higher than the value previously reported for CV measurements on *Ca*. E. gigas filaments [16], but corresponds well to the maximum value of 89×10^-18^ mol O_2_ cell^−1^ s^-1^ reported for freely moving freshwater cable bacterium filaments in microcosms as assessed by microsensor profiling [21]. Compared to unicellular aerobic sulfur oxidizers [33, 34], the cellular O_2_ reduction rate is 20-fold higher, and several orders of magnitude higher than the rate recorded for the aerobic methanotrophic bacterium *Methylovirgula thiovorans*, which is able to oxidize both methane and reduced sulfur compounds [35]. Expressed on a per biomass basis (assuming that an average *Ca*. E. gigas cell has a diameter of 3.5 µm and contains 680 fg of protein, calculated using the approach described in [21]), this translates into a biomass-specific value of up to 2700 nmol O_2_ mg protein^−1^ min^−1^. This value is 1.7-fold higher than the value reported for *Desulfovibrio termitidis* [36], which had until now the highest rate reported for microbes in the sulfur-metabolizing microbial realm [21].

Moreover, it should be noted that the O_2_ reduction rate quantified here should be considered as a conservative estimate. In our calculation we assume that all cells on the IGEs are fully functional. Yet, some cells in a filament will not be in proper contact with the gold surface of the electrode, and hence, will show no catalytic activity. Additionally, it has been shown that filaments typically contain a sizable amount of “ghost” cells, which reveal an empty cytoplasm and are considered metabolically inactive [37]. Accounting for these factors, the cellular O_2_ reduction rate of the cells that are active and electrode-connected will be higher than the value calculated. At the same time, it should also be recognized that the CV-based values recorded here represent potential rates, and as such, they illustrate the maximum capacity of the O_2_ reduction machinery of a cable bacterium cell [38]. The O_2_ reduction rate by cells residing under *in situ* conditions within the oxic zone of marine sediments is intrinsically constrained by the metabolism of the filaments (i.e. the current generated from sulfide oxidation in anoxic sediment), while CV based values are only limited by diffusion of O_2_ from the electrolyte buffer towards the sample (which is less restrictive and enables a much higher O_2_ reduction rate).

### 4.3 Different cable bacteria species have a common O_2_ reductase system

Our CV data show two types of activity by *Ca*. E. gigas filaments. One type of filaments (HRF) shows high O_2_ reduction rates, which appear to be limited by O_2_ transport towards the electrode, while a second class (LRF) shows lower rates that are not transport-limited (Figure 2). These two types of activity were also documented by Scilipoti et al. when assessing O_2_ consumption rates of freely moving cable bacteria in microcosms [21]. A small portion of filaments showed cells with high O_2_ reduction rates, while a larger cluster revealed oxic cells with lower activity [21]. Still, our data show that the LRF and HRF samples display the same apparent Michaelis-Menten constant (Figure 2, Table S2) and a similar pH dependence of the current (Figure 2), and so the mechanism of O_2_ reduction appears the same. The difference in bioelectrochemical activity between LRF and HRF filaments can be explained by a different concentration of enzymes, or by a different concentration of periplasmic redox shuttles that connect the O_2_ enzyme with the electrode surface, or by a combination of these two factors (Figure 4). Our data thus indicate that the same O_2_ reduction system is present in different species of cable bacteria (*Ca*. E. gigas and *Ca*. E. scaldis). This system is characterized by an onset potential for O_2_ reduction (*ca*. 30 mV), a limiting O_2_ concentration of 35 µM O_2_ and a *K’_m_* value of 27 µM (Table S2).

**Figure 4.**
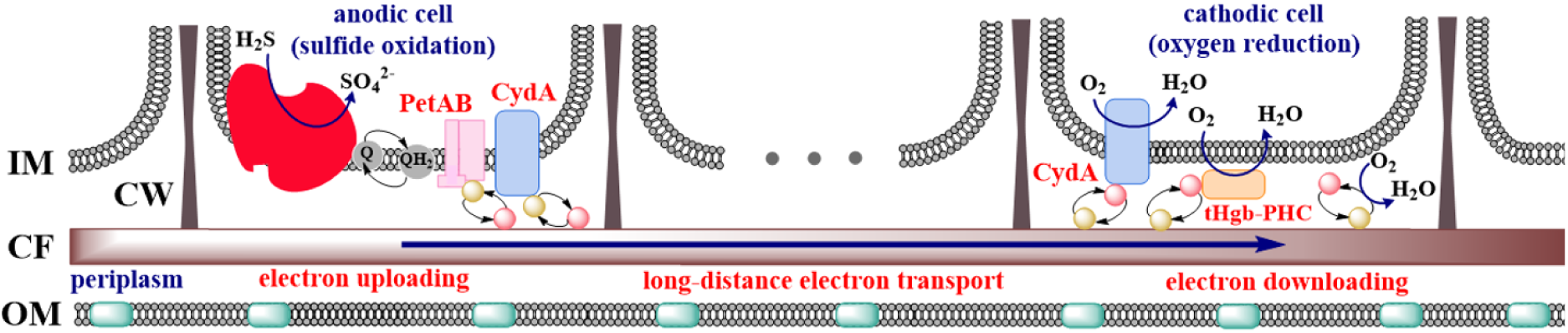
Potential electron transfer pathways for the coupling of oxygen reduction to long-range transport in *Ca*. E. gigas cable bacteria based on genomic intercomparison analysis. The anodic cells reside in deeper anoxic sediment horizons where sulfide is produced. The cathodic cells reside in a thin layer near the sediment-water interface, where O_2_ diffuses into the sediment from the overlying water column. Sulfide oxidation to sulfate is performed by an ensemble of membrane-bound and cytoplasmic protein complexes represented for simplicity by a combined structure (coloured in red). OM – outer membrane, IM – inner membrane, CF – conductive fiber, CW – cartwheel structure. Pink and yellow circles – fully oxidized and fully reduced redox shuttles, respectively; green rectangles – conducting protein.

There is a marked change in the slope of the onset potential around pH 6 (Figure 2B,E), which suggests a change in the mechanism of biological O_2_ reduction. A similar behavior has been previously seen in blue multicopper oxidases, for example, in the case of bilirubin oxidase from *Magnaporthe oryzae,* where the slope changes from −28 mV/pH to −57 mV/pH also at pH 6 [39]. An inflection in the redox potential has also been observed at near-neutral pH values for other heme-containing proteins, e.g. myoglobins [40–42], eukaryotic hemoglobins [40, 41], and peroxidases [40]. This break in the slope of the onset potential is conventionally explained by a change in the protonation state of the amino acid residues (His, Glu, Asp) ligating the active center of the metallo-proteins [39, 43, 44]. This hence suggests that proteins with metal cofactors are involved in the oxygen reduction process in cable bacteria.

### 4.4 The genetic repertoire for O_2_ reduction in cable bacteria

The recorded O_2_ reduction activity of native filaments implies that a similar O_2_ reductase system is present in different species of cable bacteria. Recently, a number of complete genomes of cable bacteria have become available [18, 26, 45, 46], and assessing the genomic repertoire of cable bacteria, there are several putative pathways for O_2_ reduction (Figure 4). All cable bacteria genomes encode a truncated hemoglobin (tHgb), which is predicted to be located in the periplasm (Figure 4) [18, 26, 45]. In some species of cable bacteria (e.g. *Ca*. E. arhusiensis and *Ca*. Electronema aureum) the tHgb gene is fused with a sequence coding for a pentaheme cytochrome c, whereas in other cable bacteria species tHgb and pentaheme cytochrome c are encoded by separate genes [18]. Nevertheless, the presence of a gene fusion in some genomes suggests that these subunits form a protein complex. In addition, certain genomes contain a second gene encoding tHgb [18]. Although little is known about the role of tHgb in bacteria, tHgb from *Bacillus subtilis* has been shown to bind and reduce O_2_ to water. Its heme redox potential shows a similar pH dependence as seen in Figure 2B,E, with an inflection point from ca. −60 to −30 mV/pH at pH 7 when tHgb is adsorbed on a graphite electrode [44] or at pH 6 when tHgb is immobilized on C_11_NH_2_-modified gold surface [47]. A similar inflection point is seen here (Figure 2, B, E). Moreover, immobilized tHgb displays a relatively high apparent catalytic rate constant of 56 s^-1^ toward O_2_ [44], which confirms that tHgb could also be involved in O_2_ reduction in cable bacteria.

All cable bacteria also contain genes encoding small periplasmic cytochromes, which are thought to enable the uploading of electrons derived from sulfide oxidation to the conductive fiber and also electron downloading from the conducting fiber towards O_2_-reducing complexes in the periplasmic space (Figure 4) [18, 45]. Although these small periplasmic cytochromes are primarily thought to enable electron transfer between different complexes (Figure 4), the involvement of periplasmic cytochromes in O_2_ reduction, as observed in *Desulfovibrio* species [48, 49], cannot be excluded based on the current dataset.

Furthermore, cable bacteria genomes encode a distant homolog of the cytochrome bd quinol oxidase subunit CydA, which is predicted to be located in the inner membrane (Figure 4). The CydA homolog in cable bacteria exhibits a distinctive domain architecture compared to the canonical CydA subunit found in other organisms, featuring a C-terminal fused cytochrome domain predicted to reside in the periplasm [18]. Notably, most cable bacteria lack the additional large subunit CydB, which is structurally near-identical to CydA, and the small subunit(s) (CydX, CydH, CydS), which comprise the full bd quinol oxidase complex [50, 51]. Complete cytochrome bd quinol oxidase complexes are present in several related anaerobic sulfate-reducing bacteria of the *Desulfobulbales* order and are known to reduce O₂ with high affinity for the detoxification of oxygen [52]. Structural analyses of the complete cytochrome bd quinol oxidase complex from *Escherichia coli*, *Geobacillus thermodenitrificans* and *Mycobacterium tuberculosis* have demonstrated that the three hemes facilitating oxygen reduction are found within the CydA subunit, with heme arrangements that vary between different species [50, 51, 53–55]. The CydB subunit is also essential for oxidase activity [56, 57]. In cable bacteria, the metabolic role of the CydA homolog remains unresolved. It has been hypothesized that CydA mediates electron transfer from the quinone pool to the periplasm [18], similar to CymA in *Shewanella oneidensis*, an inner-membrane protein which also contains a periplasmic cytochrome domain and is integral to the electron transfer process [58]. Alternatively, the periplasmic cytochrome domain of CydA might couple the electron pool in the periplasm of cells in the oxic zone to the hemes of the active site in CydA, enabling oxygen reduction. In addition, a dual functionality cannot be excluded. Experimental evidence suggests that oxygen reduction can be activated by all cable bacteria cells at all times without any delay, irrespective of their role as "cathodic" or "anodic" cells at any given time [4]. Cells can switch roles within minutes—far faster than gene expression would permit [4]. This rapid adaptation implies a constitutive expression of the genes involved in the mechanism(s) of O_2_ reduction, enabling "anodic" cells, which transfer electrons from the quinone pool to the periplasm, to immediately transform into "cathodic" cells capable of reducing oxygen via the periplasmic electron pool. In addition to CydA, cable bacteria encode a Rieske Fe-S protein (PetA) and a membrane-bound cytochrome b-domain protein (PetB), which have been proposed to be involved in electron transfer from the quinone pool to the periplasm (Figure 4) [18]. The CydA homolog could serve as a flexible complex, serving both an electron transfer from the quinone pool to the periplasm and an oxygen reductase linked to the periplasmic electron pool, although this requires further experimental validation.

Besides these mechanisms present in all cable bacteria, there are genetic indications for terminal O_2_ reductases that are only present in specific species. Although absent in most cable bacteria, the recently discovered *Ca*. E. antwerpensis contains a full cytochrome *bd* quinol oxidase complex (CydABX) [26], in addition to a CydA homolog. Canonical CydABX complexes receive electrons from reduced quinones in the membrane for the O_2_ reduction. Another example is found in the genomes of *Ca*. E. scaldis, *Ca*. E. communis and *Ca*. E. laxa [18, 26, 45], which code for the cytochrome *c* aa3-type oxidase complex CoxABCD. This complex could receive electrons from periplasmic cytochromes (Figure 4), instead of from cytochrome c reduced by canonical complex III as this complex is missing in cable bacteria [18]. Nonetheless, based on the similar high ORR between *Ca*. E. scaldis and *Ca*. E. gigas, as presented in Figure 1 and Table S2, the presence of cytochrome c oxidase has a limited impact on the potential ORR activity.

### 4.5 A model for O_2_ reduction coupled to long-range electron transport

Our data provides further insight into how O_2_ reduction is coupled to the long-range electron transport in the fiber network of cable bacteria. The O_2_ reductases are most likely not in direct contact with the electrode, and so the observed capability of O_2_ reduction requires a suitable transport path for the electrons from the electrodes to the O_2_ reductases embedded in a filament. We conjecture that soluble periplasmic c-type cytochromes may act as shuttles between electrodes and O_2_ reductases (Figure 4). The presence of such cytochromes is demonstrated in the genomes of cable bacteria [18, 26, 45], while Raman spectroscopy has revealed a high content of cytochromes in native cable bacteria, but a near absence in fiber skeletons [9, 14], thus indicating that the periplasm is enriched in extractable (i.e. soluble and putatively motile) cytochromes.

The proposition of cytochrome-mediated electron transfer in O_2_ reduction is supported by the absence of any O_2_ reduction activity during cyclic voltammetry when the cable bacteria were treated with N-acetylmethionine (Figure S4), which deactivates c-type cytochromes [24, 25]. The involvement of cytochromes was additionally confirmed by our electrochemical gating results, which revealed a potential of ∼ −100 mV for the redox species involved in conduction (Figure 3). This value is close to the half-wave potential as recorded in the CVs of native filaments (Figure 1D,E,H), which suggests that cytochromes likely act as redox shuttles in the oxygen reduction process [59].

The proposition that periplasmic c-type cytochromes play a crucial role in mediating electron transport through the periplasm explains the electron flow observed under the different experimental conditions examined here (Figure 5). Under *in situ* conditions, the cytochromes shuttle electrons between the conductive fiber and oxygen-reducing complexes, which either perform sulfide oxidation or oxygen reduction [10] (Figure 5A). Likewise, in our cyclic voltammetry experiments, the cytochromes are shuttling electrons from the electrode surface to the oxygen reduction complexes (Figure 5B). In the electrochemical gating experiment, the cytochromes are shuttling electrons from the electrode to the conductive fiber as well as the oxygen reduction complexes (Figure 5C).

**Figure 5.**
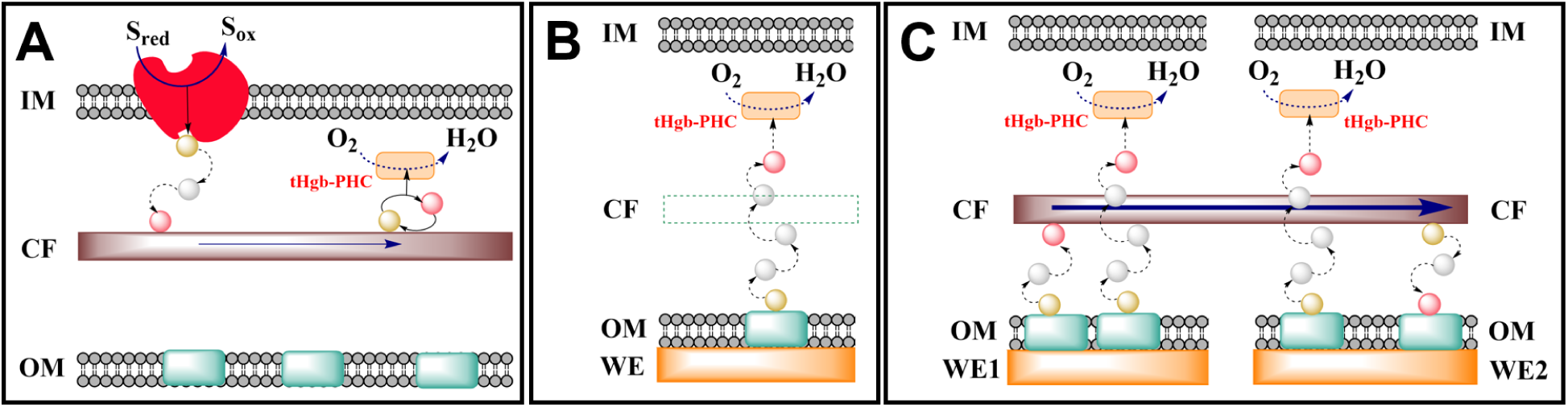
Schematic overview of possible electron transfer pathways in cable bacteria under different experimental conditions investigated here. This is a simplified version of Figure 4 showing only a tHgb-PHC. (A) Electron flow under natural conditions. (B) Electron flow during voltammetry (C) Electron flow during electrochemical gating conditions. OM – outer membrane, IM – inner membrane, WE – working electrode (WE1 and WE2 – source and drain electrodes (see Fig. 1A)), CF – conductive fiber. Pink, yellow and gray circles – fully oxidized, fully reduced and state-altering redox shuttles, respectively; red shape – sulfur-oxidizing enzymes, blue crescents – inner-membrane oxygen-reducing proteins, green rectangles – conducting protein.

In all cases, the periplasmic c-type cytochromes enable an efficient electron transport through the periplasm, which allows to sustain high cellular O_2_ reduction rates, both under *in situ* conditions as when performing CV experiments (Figure 5). In the latter experiments, the conductive fibers do not play a role in the electric currents recorded. This is confirmed by the electrochemical gating experiment, in which the source-sink current vanished at higher oxygen concentrations (Figure 3D), thus signifying a deterioration of the fiber conductivity. Yet, under these same aerobic conditions, cyclic voltammetry revealed that the cells were still capable of O_2_ reduction activity, and so redox shuttles must remain functional. The degradation of the fiber conductance by oxygen could be potentially caused by an irreversible oxidation of the Ni/S supramolecular structure that is embedded within the fibers [14, 15]. Note that the disappearance of the source-drain current at high O_2_ concentrations (Figure 3D) also implies that the redox shuttles cannot be responsible for any electron transfer along the longitudinal axis of a filament, since these redox shuttles remain functional when O_2_ is present.

Previously, it has been proposed that cable bacterium filaments could be potentially used as oxygen-reducing biocathodes in electrochemical applications [19]. In our experiments, the maximum current density recorded was ∼ 4 A m^-2^, assuming that the IGEs were covered with a single layer of cable filaments (no superposition of filaments). This is a high current output for a 2D surface under diffusion limitation, and comparable to the highest current densities obtained for oxygen-reducing biocathodes with extended 3D structure and improved gas-liquid oxygen transfer [60, 61]. However, despite the exceptionally high current density, the relatively low onset potential and the limited operational stability under aerobic conditions make the practical implementation of CB-based biocathodes challenging.

## CONCLUSIONS

In this study, we combined cyclic voltammetry and electrochemical gating to elucidate how O_2_ reduction in cable bacteria is linked to the long-distance electron transport along the conductive fiber network. Our data show that the fibers themselves only perform longitudinal electron transport, and are not electrochemically active towards oxygen. In contrast, native cable bacterium filaments are capable of high oxygen reduction rates, thus demonstrating that dedicated enzyme systems in the periplasm or inner membrane are responsible for the observed O_2_ reduction. Together, our data provide empirical support to a model in which periplasmic c-type cytochromes play a crucial role in mediating electron transport through the periplasm, shuttling electrons between dedicated respiratory complexes and the conductive fiber network. As such, this study resolves another crucial piece of the unique electrogenic metabolism that fuels long-distance electron transport in cable bacteria, while clarifying the application potential of cable bacteria-derived enzyme systems in BES (Bio-electrochemical Systems) technologies.

## ACKNOWLEDGMENTS

The research is financially supported by Research Foundation Flanders (FWO SBO grant S004523N, FWO project G0ADR25N and FWO PhD fellowship 11D7822N) and by the University of Antwerp via the TopBOF program. FJRM received additional support from the Horizon Europe research and innovation programme EIC Pathfinder under grant agreement PRINGLE 101046719.

## TABLE OF CONTENTS

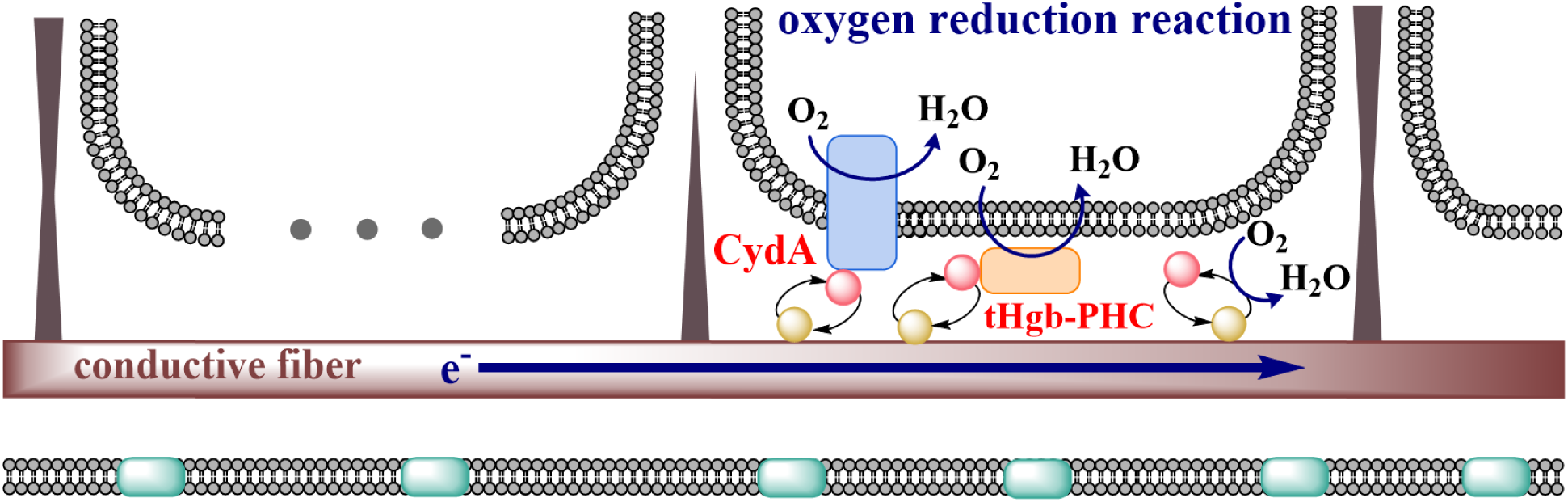

