## Supplementary figures and tables for "Highly efficient bio-catalytic oxygen reduction coupled to long-range electron transport in cable bacteria"

### SUPPORTING INFORMATION

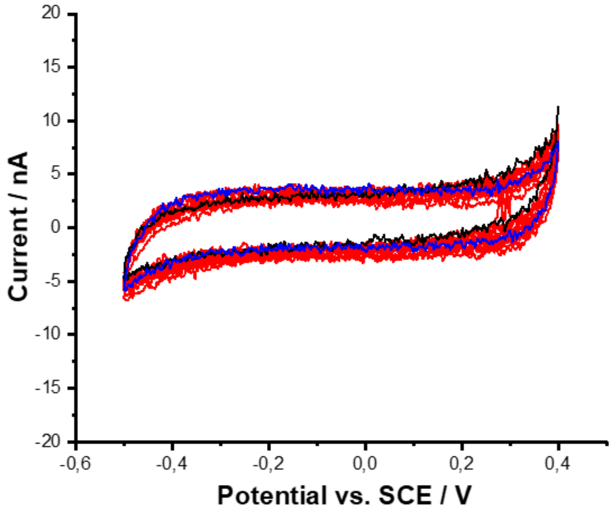


**Figure S1.** Control experiment for Cyclic Voltammetry. Voltammograms of a non-biomodified thiolated gold electrode at different concentrations of $O_2$ (0-58 μM) in 50 mM potassium phosphate buffer (pH 7.0). Scan rate: 20 mV $s^{-1}$. No $O_2$ reduction activity is apparent.

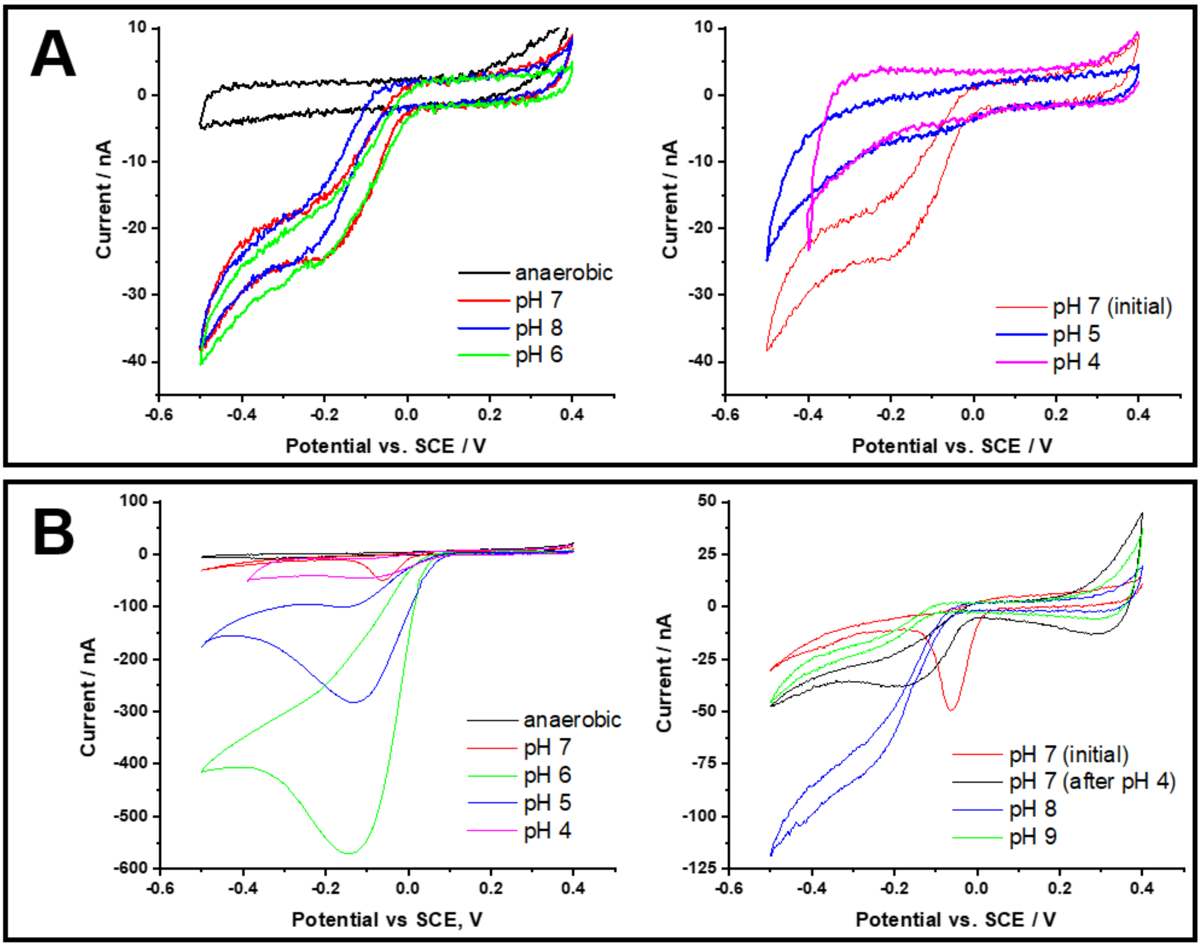


**Figure S2.** Representative CVs of low-rate (A) and high-rate filaments (B) recorded in electrolyte buffer solutions at constant 12.5 μM $O_2$ but different pH values. Different buffer solutions (50 mM sodium acetate, 50 mM potassium phosphate and 50 mM sodium bicarbonate) were used to adjust pH values to 4-5, 6-8 and 9, respectively.

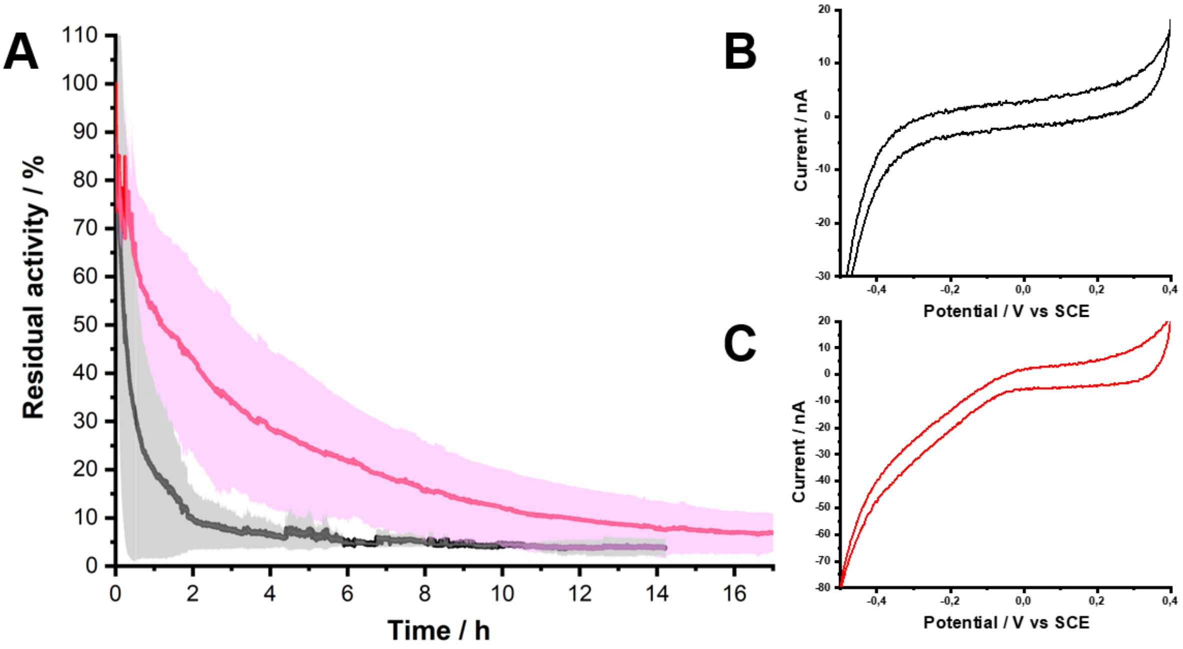


**Figure S3.** A) Temporal evolution of the cathodic current measured amperometrically at a fixed -0.24 V vs SCE in an air-saturated buffer solution (50 mM potassium phosphate, pH 7.0) for low-rate filaments (black curve) and high-rate filaments (red curve). B) CVs recorded for the low-rate filaments (after the stability test. C) CVs recorded for high-rate filaments (C) after the stability test.

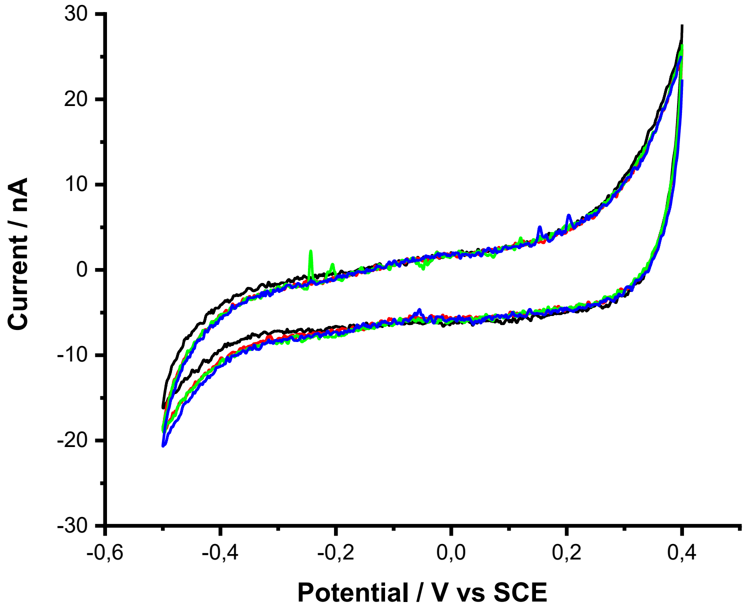


**Figure S4.** Representative CVs of intact *Ca*. E. gigas treated with 100 mM N-acetyl-L-methionine recorded in buffer solution (50 mM potassium phosphate; pH 7.0) at different concentrations of $O_2$: 13 µM (black), 24 µM (red), 36 µM (green), 46 µM (blue). Potential scan rate: 20 mV $s^{-1}$. No $O_2$ reduction activity is apparent.

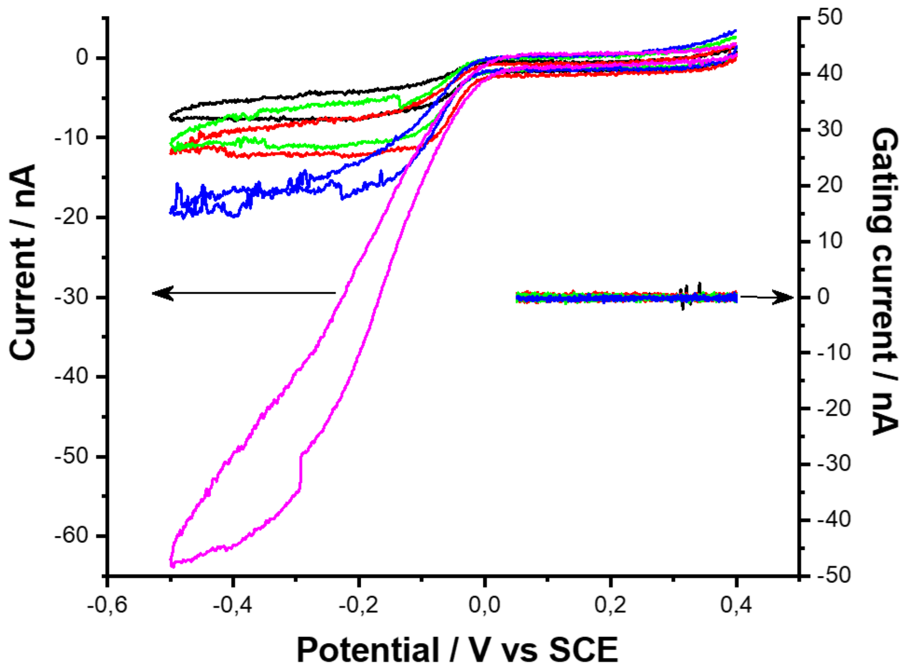


**Figure S5.** Electrochemical gating experiments. Left axis: CVs of the source electrode at different $O_2$ concentrations: 0 µM (black), 3.2 µM (green), 9.5 µM (red), 21.8 µM (blue), and air-saturated (magenta). Right axis: dependence of the source-drain current on the potential of the source electrode for intact cable bacteria in oxygen-saturated conditions (50 mM potassium phosphate buffer solution; pH 7.0). Source-drain voltage: 0 mV (black curves), 3 mV (red curves), 6 mV (green curves), and 10 mV (blue curves). Scan rate: 5 mV $s^{-1}$.

**Table S1**. Calculation of cellular $O_2$ reduction rates. Data are tabulated for the oxygen reduction rate of LRF and HRF samples deposited on the thiolated gold surface. Characteristics of the HRF are marked in bold font and grey. The maximum current is derived from CVs. An average cell length of 3 µm was used to calculate the number of cells from the electrochemically active length (as determined by light microscopy of the IGE).

| Sample | Electrochemically active length, µm | Number of cells | Maximum current, nA | Electron flow, electrons $cell^{-1}$ $s^{-1}$ $\times 10^{-7}$ | Cellular $O_2$ reduction rate | |
|---|---|---|---|---|---|---|
| | | | | | molecules $O_2$ $cell^{-1}$ $s^{-1}$ $\times 10^{-6}$ | mol $O_2$ $cell^{-1}$ $s^{-1}$ $\times 10^{18}$ |
| **1** | **21105** | **7035** | **125.8** | **11.2** | **27.9** | **46** |
| 2 | 9604 | 3201 | 19.0 | 3.7 | 9.3 | 15 |
| 3 | 23833 | 7944 | 42.1 | 3.3 | 8.3 | 14 |
| 4 | 36391 | 12130 | 106.3 | 5.5 | 13.7 | 23 |
| **5** | **18008** | **6003** | **300.1** | **31.2** | **78.0** | **130** |
| 6 | 16010 | 5337 | 38.0 | 4.4 | 11.1 | 18 |
| **7** | **38706** | **12902** | **164.2** | **7.9** | **19.9** | **33** |

**Table S2**. Michaelis–Menten kinetic parameters obtained for the LRF and HRF (bold font and grey background) filaments of *Ca*. Electrothrix gigas and for *Ca.* Electrothrix scaldis GW3-3.

| **Sample** | **Cable bacteria species** | **Type** | $K'_m$, **µM** | $I_{max}$, **nA** |
|---|---|---|---|---|
| 1 | *Ca*. E. gigas | LRF | 33.8 | 64.9 |
| 2 | *Ca*. E. gigas | LRF | 25.8 | 65.6 |
| 3 | *Ca*. E. gigas | LRF | 23.9 | 29.3 |
| 4 | *Ca*. E. gigas | LRF | 26.6 | 4.8 |
| **5** | ***Ca*. E. gigas** | **HRF** | **14.2** | **138.9** |
| **6** | ***Ca*. E. gigas** | **HRF** | **43.8** | **193.9** |
| **7** | ***Ca*. E. gigas** | **HRF** | **24.6** | **183.1** |
| *8* | *Ca*. E. scaldis GW3-3 | LRF | 31.6 | 188.8 |
| *9* | *Ca*. E. scaldis GW3-3 | LRF | 27.2 | 169.1 |
